# Cellular Context Influences Kinase Inhibitor Selectivity

**DOI:** 10.1101/2025.10.09.681452

**Authors:** Matthew J. Binder, Frances M. Bashore, Kaitlin K. Dunn Hoffman, Cameron Damgaard, Michael Slater, David H. Drewry, Matthew B. Robers, Alison D. Axtman

## Abstract

A pivotal part of kinase chemical probe and drug development is assessment of the selectivity of a putative lead compound. While there is no consensus around the panel size or the type of assay(s) that are most appropriate, there is concurrence that gauging the number of on- and off-targets of a kinase inhibitor is essential. As pharmacology takes place in cells, we have compared profiling results for ten kinase inhibitors generated using the cell-free assays to those obtained when a panel of cellular target engagement NanoBRET assays is used to assess selectivity in intact cells. This is the first systematic comparison of these two approaches across a broad kinase panel. Comparison of the data sets demonstrates divergent results that can influence chemical probe prioritization. We identify unanticipated kinase interactions in cells for type II kinase inhibitors that are not observed in biochemical, cell-free systems. Furthermore, we characterize TPKI-39 as a DDR1, DDR2, and FLT1 chemical probe based on its in-cell selectivity profile.

**For Table of Contents Only:** 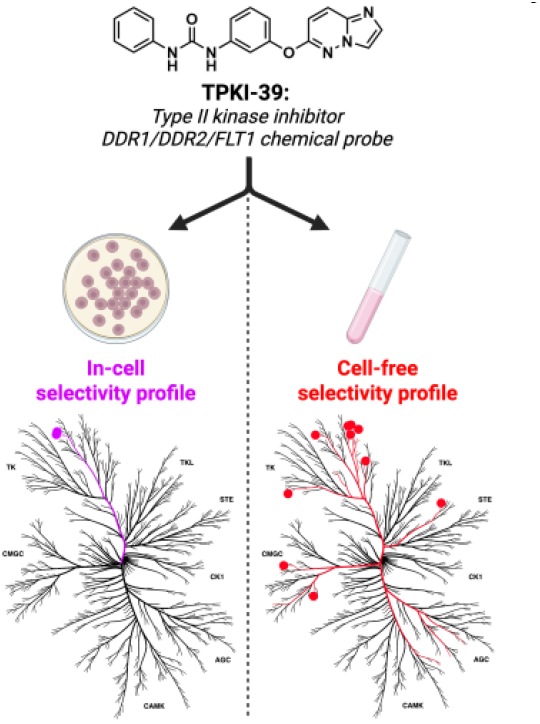

## Introduction

Small molecule chemical probes are essential tools that enable the study of biological pathways and disease.^1-3^ By identifying and utilizing potent and selective inhibitors, termed chemical probes, the function of proteins and their downstream biological effects, especially implications in disease phenotypes, can be illuminated. Chemical probes are distinct from drugs in that the focus of their development centers around potency and selectivity rather than safety and efficacy. First suggested by Workman and Collins in 2010, a set of criteria have been agreed upon by the Chemical Probes Portal and Structural Genomics Consortium (SGC) defining chemical probes.^4^ These criteria include biochemical potency <100 nM, in-cell potency <1 µM, >30-fold selectivity over closely related proteins, and availability of a structurally related negative control compound.^1, 2^

Selectivity is a key metric for chemical probes, perhaps especially so for those targeting protein kinases. Protein kinases are a class of enzymes that catalyze transfer of the γ-phosphate group of ATP onto serine, threonine, or tyrosine substrate residues. As all kinases must bind ATP to carry out their enzymatic activity, protein kinase domains are comprised of certain canonical structural features that can make achieving sufficient selectivity between on- and off-target ATP sites challenging. Kinases can be further classified as in an “active” or “inactive” conformation based on the relative orientations of these structural features.^5^ Namely, these include the αC-helix and DFG (Asp-Phe-Gly) motif of the activation segment.

The typical kinase domain is divided into an N-terminal lobe and larger C-terminal lobe, connected by a hinge region. This hinge region serves as the interface between the two lobes and is the orthosteric binding site for both ATP and ATP-competitive kinase inhibitors.^6^ The N-terminal lobe contains the αC-helix, which makes a critical salt bridge interaction with the β3-sheet in the active conformation. This positioning, where the helix is pointed inward, is referred to as the “αC-in” orientation. The DFG motif, contained within the C-terminal lobe, begins the activation loop of the kinase. The positioning of the aspartic acid residue of the DFG motif is another key determinant of kinase activity. In the active state, the Asp is directed inward (DFG-in) and can coordinate Mg^2+^, which is important for ATP binding.

Certain small molecule inhibitors have been found to bind exclusively to either the active or inactive conformation of their target kinases and have been classified based on these preferences. As many as seven different kinase inhibitor types have been proposed, including both orthosteric and allosteric modulators, as well as covalent binders.^7^ Here, we will focus on the most common classes of orthosteric inhibitors: type I and type II. Type I inhibitors bind to the ATP-binding site of the active kinase conformation, with both the αC-helix and DFG motif positioned inward. Type II inhibitors, conversely, bind the ATP-binding site of the inactive kinase, with αC-out and DFG-out. This structural difference can be seen in Figure 1, which compares c-Abl bound to dasatinib (left)^8^ and imatinib (right),^9^ type I and type II inhibitors, respectively. Originally, it was proposed that type II inhibitors may be inherently more selective than type I due to the more unique conformational states of inactive kinases, increasing their value for chemical probe and drug discovery efforts.^10^ However, further investigation has largely disproven this idea.^11^ In fact, type II inhibitors may sample different conformational space that imparts them with unique selectivity profiles when compared to their type I counterparts.^11, 12^ Regardless, selectivity remains a top priority when developing compounds.

**Figure 1.**
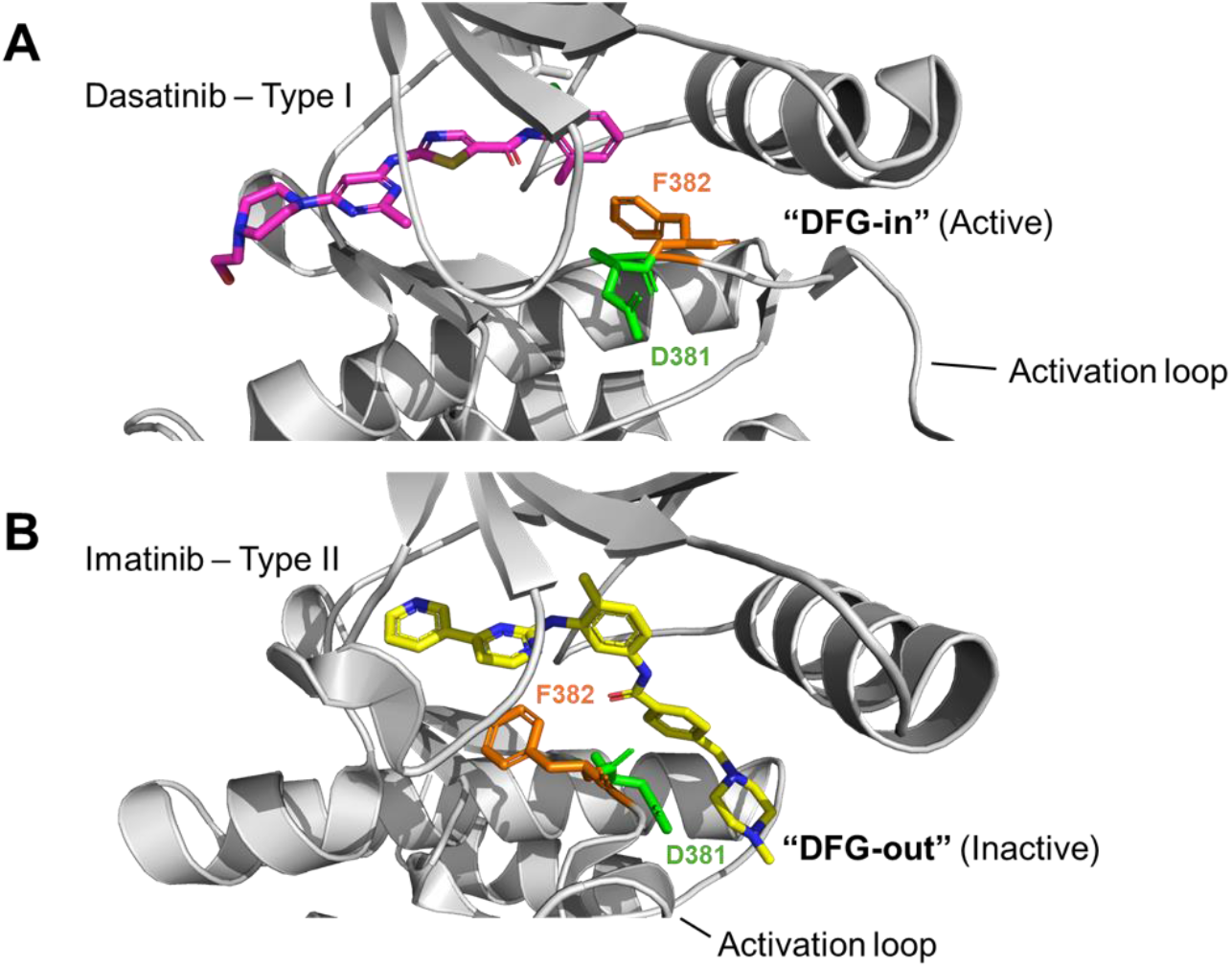
Structural comparison of type I versus type II binding modes. (A) Crystal structure of dasatinib (pink), a type I inhibitor, bound to c-Abl (PDB code: 2GQG). (B) Crystal structure of imatinib (yellow), a type II inhibitor, bound to c-Abl (PDB code: 1IEP). The activation loop containing the DFG motif is highlighted.

To quickly and comprehensively evaluate the selectivity of a lead inhibitor, high-throughput kinase profiling panels have been developed. These panels contain representative kinases from the major families:^13^ TK, TKL, AGC, CAMK, CK1, CMGC, STE, and atypical, as well as mutant kinase constructs and lipid kinases. To date, the Eurofins DiscoverX panel has achieved the broadest coverage, with 468 kinases included, representing more than 80% of the human kinome. This panel consists of competitive binding assays between a promiscuous kinase ligand attached to a solid support and a compound of interest.^14^ DNA-tagged kinases are allowed to bind to the promiscuous ligand and compound is added at a single concentration. While off-target hits are displaced from the solid support by the compound of interest, kinases that remain bound after washing can be quantified via qPCR. Results are returned as percent of control (PoC), which is relative to the assay result when no competing compound was added. This panel allows for the quantitative evaluation of kinome-wide selectivity in a cell-free format, providing a convenient and robust platform to identify novel interactions and chemical starting points for understudied kinases.

For a complementary evaluation of target occupancy in a physiological context, NanoLuc-bioluminescence resonance energy transfer (NanoBRET) cellular target engagement assays can be performed in live cells. NanoBRET assays, developed by Promega in collaboration with the Structural Genomics Consortium, allow for a direct assessment of the engagement of a protein target by a compound in a cellular setting.^15-17^ These assays require a tracer molecule, a heterobifunctional compound consisting of a ligand with known affinity to the target protein conjugated to a fluorophore. This tracer molecule binds to the transiently transfected NanoLuc (NLuc)-tagged full-length kinase, and upon the addition of an NLuc substrate, an energy transfer occurs between NLuc and the fluorophore on the tracer. Addition of a compound that competes with binding of the tracer results in a loss of luminescence signal. Akin to the biochemical kinome profiling techniques, NanoBRET assays run in parallel can also be used to quickly evaluate selectivity of a compound across the kinome.^18^ To date, NanoBRET assays have been enabled for more than 300 kinases. Kinase targets with NanoBRET assays are well-distributed among the kinase families and generally overlap with the biochemical panels.^19^ By screening a compound of interest at a single concentration, percent occupancy can be determined for each kinase in the panel. The NanoBRET assay assessment of direct target engagement contrasts with immunoassays, which indirectly confirm biological effect by assaying downstream kinase substrates of the kinase of interest. Herein, we systematically compare kinase inhibitor selectivity obtained through cell-free and in-cell profiling approaches and highlight the power of using in-cell target engagement assays to identify previously unknown kinase targets and uncover the exquisite cellular selectivity of certain type II inhibitors. We demonstrate that cellular profiling of kinase inhibitor selectivity is informative for the development of high-quality chemical probes. The implications of our findings should resonate with scientists across multiple fields with interest in optimization of kinase-targeting small molecules via the most rigorous methodology.

## Results and Discussion

Comparison of type I and type II inhibitors in larger cell-free assay panels has taught us that there are no universal trends that establish one or the other binding mode as a more selective class.^20^ A side-by-side comparison of global cell-free and in-cell data has not been reported in the literature, providing an opportunity to benchmark and explore the value of using both methods for chemical probe optimization.

### Profiling of the binding of widely used kinase inhibitors surfaces otherwise undetected in-cell interactions of type II inhibitors

Taking advantage of previously published and/or database-housed cell-free datasets, we planned experiments aimed at comparing the cell-free broad profiling results with the results obtained when in-cell profiling was executed. The Library of Integrated Network-based Cellular Signatures (LINCS) houses a compendium of cell-free datasets.^21^ These data, alongside published profiling efforts, enable the designed comparisons.^14, 22, 23^

#### Type II kinase inhibitors sample unique intracellular conformational space

To start, a sentinel set of four widely-used kinase inhibitors, dasatinib, sorafenib, AST487, and foretinib, (Figure 2A) were chosen for which there was published broad biochemical profiling data as well as dose– response follow-up values. This included one type I and three type II inhibitors with varied kinome-wide selectivity. Based on their cell-free profiling,^14^ these inhibitors are incredibly promiscuous, allowing us to sample many putative inhibitor-kinase interactions. The promiscuity of these kinases can be better understood if the number of wild-type kinases that bind with high affinity (K_d_ <100 nM) in the Eurofins DiscoverX cell-free panel are tabulated: 52 for dasatinib, 16 for sorafenib, 66 for AST487, and 111 for foretinib. Furthermore, it was anticipated that through use of full-length kinases in their dynamic intracellular state in the NanoBRET assay, novel interactions might also be pinpointed. These inhibitors were screened in the cell-free assay panel with the broadest kinome-wide coverage, which is offered by Eurofins DiscoverX.^14^ This panel employs binding assays that are executed in the absence of ATP, often with truncated kinases. In the 468-member *scan*MAX panel, for example, 78% of kinases (363) are partial constructs while 22% of kinases (105) are full-length. We profiled these four inhibitors using a cell-based NanoBRET selectivity panel consisting of 240–300 kinases. To compare the results generated in the cell-free platform with those produced using the cell-based selectivity screen, we examined the data obtained after running the binding assays in dose–response format (K_d_ values for the Eurofins DiscoverX panel^14^ and IC_50_ values for the NanoBRET panel) for human, wild-type kinases. This analysis is shown in Figure 2, where all values between 1 and 10000 nM are plotted. A correlation line has been drawn to better illustrate those kinases that bind with equal affinity in cell-free and cellular assays. Kinases that diverge from the central line are either more active in cell-free assays (blue) or in cell-based assays (red and green). We observed consensus in the binding of the inhibitor to many of the same kinases in each panel. It is worth noting that the primary kinase targets of these inhibitors, when present in both panels, were identified as high affinity binders. As an example, for dasatinib, which inhibits BCR-ABL, EGFR, SRC, LCK, YES, FYN, KIT, EPHA2, and PDGFRB,^24^ high affinity binding to ABL, SRC, LCK, YES1, FYN, KIT, and EPHA2 was identified in both the in-cell and cell-free data. For all these kinases, however, the cell-free affinity was <1 nM so they are not amongst the kinases plotted in Figure 2C. For sorafenib, RET, FLT3, and KIT are primary target kinases^24^ found to bind with high affinity in both the in-cell and cell-free panels that are also plotted in Figure 2D. Other primary targets, including B/C-RAF, VEGFR1/2/3, and PDGFRB,^24^ were absent from either one or both panels. Finding these primary drug targets in our data sets further validates our method.

**Figure 2.**
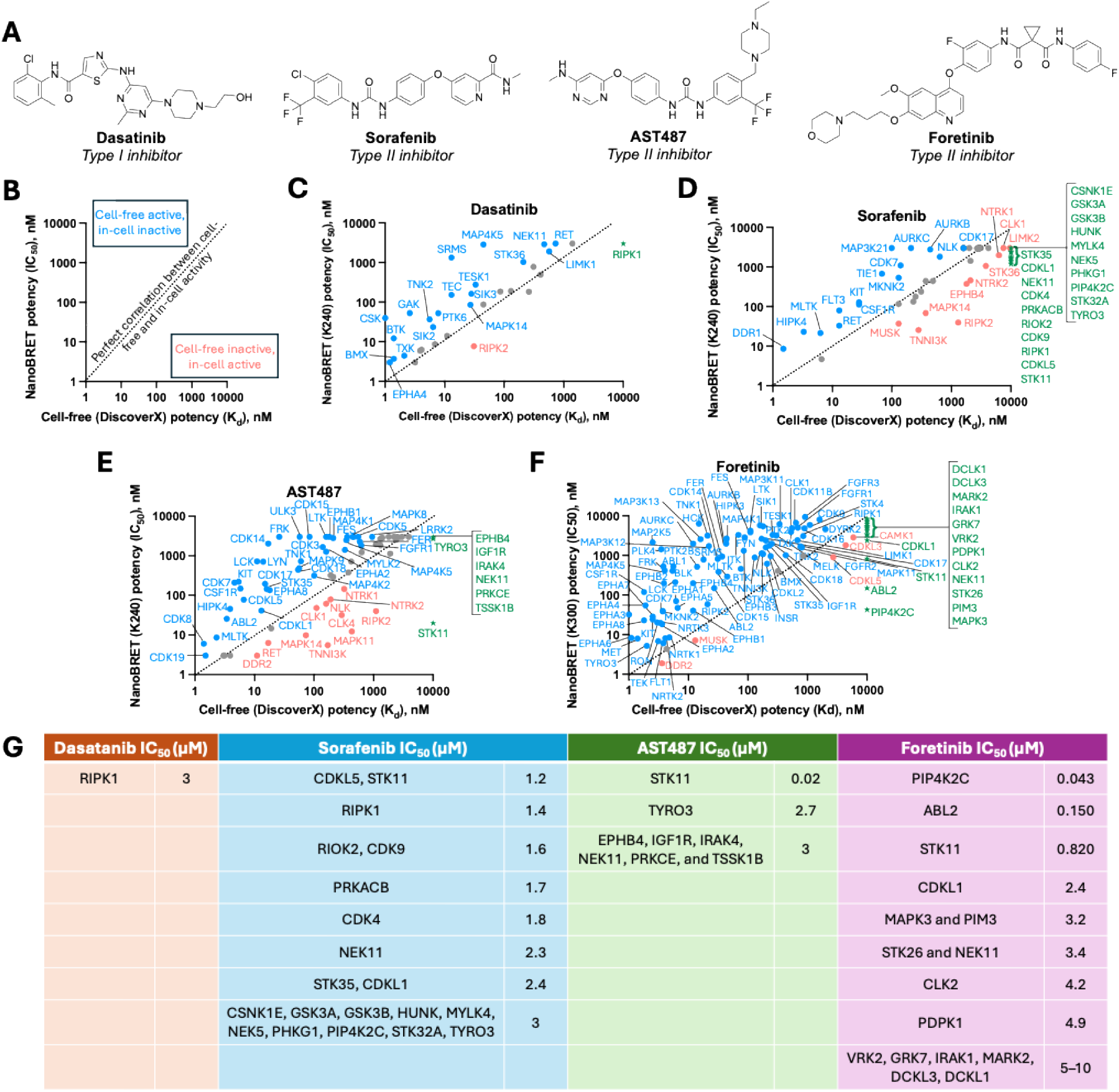
Comparison of K_d_ and IC_50_ values generated for select kinase inhibitors in cell-free binding and cell-based binding assays, respectively, when inhibitors were screened in dose–response format. (A) Structures of a sentinel set of non-selective kinase inhibitors chosen for in-cell selectivity assessment. (B) A key is included to aid in data interpretation. Data is shown for (C) dasatinib (type I), (D) sorafenib (type II), and (E) AST487 (type II), and (F) foretinib (type II). (G) A table is included to highlight those kinases present in both panels that were active in the cell-based, but not cell-free assay panel. Kinases colored green in the panels C–F are those featured in the table (G), for which no binding was observed in the cell-free assay but an IC_50_ was recorded in the corresponding cell-based assay.

For each inhibitor, the cell-based selectivity panel identified fewer kinases that bound with high affinity than in the cell-free panel. This is likely the result of poor cellular permeability of the inhibitors coupled with ATP competition in cells, as ATP is not present in the cell-free DiscoverX panel assays. Unsurprisingly, more kinase–inhibitor interactions overall were identified when the larger in-cell panel of 300 kinases was used to profile foretinib (Figure 2F) versus when only 240 kinases were included in the panel (Figure 2C–E). The number of interactions could also be influenced by the greater promiscuity of foretinib versus the other inhibitors. The four inhibitors consistently bound with higher affinity in the following NanoBRET assays: DDR2, RIPK2, CLK1, NTRK1, NTRK2, MAPK14, MUSK, and TNNI3K. A table of interactions captured in the cell-based assay that were missed in cell-free assays are captured in Figure 2G. These interactions are shown in Figure 2C–F as green stars and text. Notably, potent in-cell affinity of AST487 for STK11 was observed. Strikingly, many more cell-based interactions were identified for the type II inhibitors when compared to the type I inhibitor, revealing binding interactions that may have been otherwise overlooked. The number of ‘missed interactions’ is similarly increased when comparing type II to type I inhibitors. It is possible that many kinases may adopt DFG-out conformations that are not sampled with active protein in cell-free assays. In-cell profiling of type II inhibitors, therefore, captures additional interactions when compared to their cell-free selectivity profile.

#### Cell-free and in-cell selectivity fingerprints confirm unique type II trends

To expand on this preliminary analysis, we next compared single-concentration selectivity data for two type II inhibitors, LXH254^22^ and ponatinib, and a type I kinase inhibitor, AMG925,^23^ in the same panels of binding assays (Figure 3A).^25^ For this analysis, the single concentration (1 µM) data from the cell-free profiling (DiscoverX) was plotted versus the percent occupancy generated in the cell-based NanoBRET selectivity panel consisting of 240–300 kinases. DiscoverX percent of control values were converted to percent inhibition values to simplify data display (% inhibition = 100 - PoC). Only the wild-type, human kinases that demonstrated >50% binding in both assay formats are plotted in Figure 3. Once again, a correlation line was drawn to help with data visualization. In-cell dose–response follow-up for LXH254 demonstrated the following IC_50_ values: DDR1 IC_50_ = 25 nM, DDR2 IC_50_ = 86 nM, and MAPK11 IC_50_ = 870 nM. This finding, where percent occupancy data is predictive of dose–response follow-up results, has previously been observed for inhibitors of CK2 and PIKfyve.^26-28^ Corresponding K_d_ values for LXH254 were reported: DDR1 K_d_ = 1.8 nM and DDR2 K_d_ = 10 nM.^14^ The few kinases plotted in Figure 3C is unsurprising given the high selectivity of LXH254.

**Figure 3.**
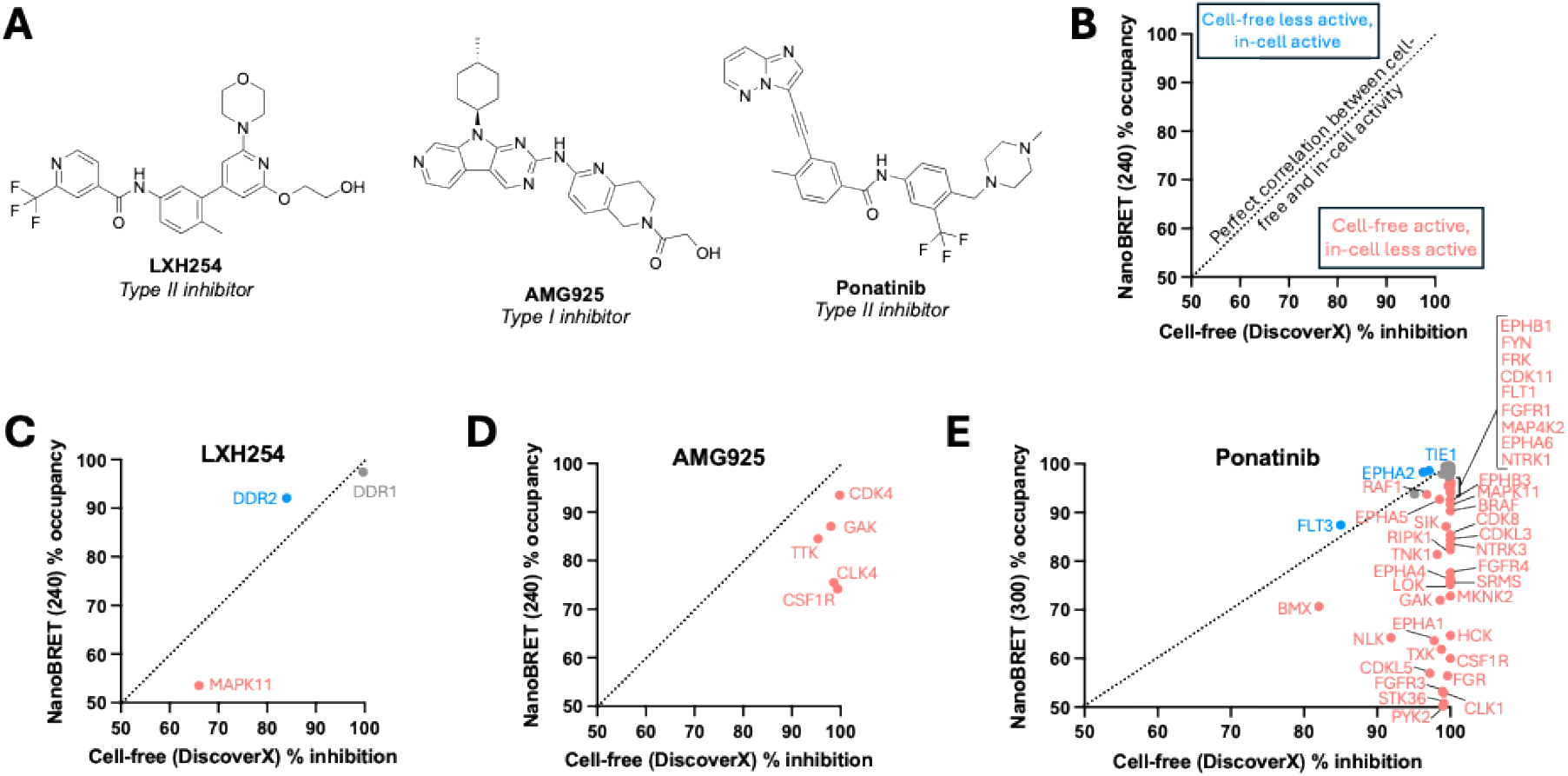
Comparison of percent inhibition and percent occupancy values generated for two kinase inhibitors in cell-free binding and cell-based binding assays, respectively, when inhibitors were profiled at 1 µM. (A) Structures of all compounds shown in panel (A). (B) A key is included to aid in data interpretation. Data is shown for (C) LXH254 (type II), (D) AMG925 (type I), and (E) ponatinib (type II). Calculation of % inhibition: 100 – PoC.

Several of the kinases with highest cell-free affinity^23^ were also found to have high affinity in cells for AMG925 (Figure 3D). When the same cell-free binding assays were executed in dose– response format to determine K_d_ values, high affinity binding of AMG925 was confirmed for many of the same kinases. This finding validated the single concentration cell-free screening as predictive of the more comprehensive dose–response profiling.^23^ The in-cell data, however, suggests that the potency of many of these kinases is diminished in intact cells, and the in-cell selectivity of AMG925 is much improved when compared to its cell-free profile. Only 30 kinases bind AMG925 with high affinity (<10 PoC) at 1 µM in the cell-free panel. As is often the case, even fewer kinases (6 with % occupancy >50) bind when AMG925 is profiled in cells. As previously discussed, this is due to ATP competition and compound permeability being factors in the in-cell assays that are absent when the cell-free system is used. Kinases that bind to AMG925 with high affinity in the cell-free panel (>99% PoC) that are also present in the in-cell panel, including CSNK2A2, NTRK2, DYRK1A, FLT3, CLK1, and DYRK1B, still bind to AMG925 in cells but with an occupancy value <50% (8–39% occupancy).

Since ponatinib was screened in a larger cell-based panel, more overlapping kinases were identified in common between the cell-free and cell-based profiling. When compared to LXH254 and AMG925, ponatinib is much more non-selective, binding with >50% occupancy to 63 of 300 kinases in the cell-based panel and <50% of control for 253 of 403 wild-type human kinases in the cell-free panel. The overlap of these kinases was high, resulting in many points plotted in Figure 3E. Only three kinases showed higher in-cell affinity when compared to cell-free binding data: FLT3, EPHA2, and TIE1. Once again, STK11 emerged as demonstrating high affinity binding in the NanoBRET panel (84% occupancy) but weak inhibition in the cell-free binding assay (28% inhibition), solidifying it as an outlier kinase with unique in-cell binding characteristics. These data support that even in single-concentration profiling, in-cell profiling of type II inhibitors can identify distinct interactions that are not captured in cell-free screening.

### Frequent hitter kinases revealed by live-cell NanoBRET target engagement

Intrigued by the novel interactions identified for type II inhibitors in our sentinel panel (Figure 2A: sorafenib, AST487, and foretinib) in kinome-wide NanoBRET assays, we explored whether a subset of kinases may represent “frequent hitters”^29^ and demonstrate intrinsic vulnerability to engagement by this chemotype in live cells. Therefore, beyond the sentinel set, we profiled a broader panel of FDA-approved type II kinase inhibitors^24^ and repeatedly detected robust cellular engagement of three kinases—PIP4K2C, STK11 (LKB1), and RIPK1—that are not cataloged as canonical type II kinase targets (Figure 4, Figure S1).^11^ Their appearance across multiple chemotypes implies that each can adopt a type II/DFG-out back pocket pose, expanding the structural definition of the type II kinome. From a discovery standpoint, compounds that exploit the type II binding channel may represent attractive starting points for chemical probe campaigns aimed at these three kinases. From a translational perspective, their hidden engagement by marketed drugs adds mechanistic context to both efficacy and safety observations in the current targeted kinase inhibition landscape. There are limited examples of potent and/or selective type I kinase inhibitors of PIP4K2C (a lipid kinase), STK11 (a tumor suppressor kinase), and RIPK1 (an innate immunity kinase). RIPK1 has also been targeted with type III allosteric inhibitors that extend into the back pocket of the kinase.^30-33^ Several approved type II kinase inhibitors reach free plasma or tissue concentrations that overlap or hover near their NanoBRET potencies for these kinases. Understanding such interactions helps rationalize unexpected clinical activity (benefit) or adverse events (liability) and guides biomarker selection for combination studies.

**Figure 4.**
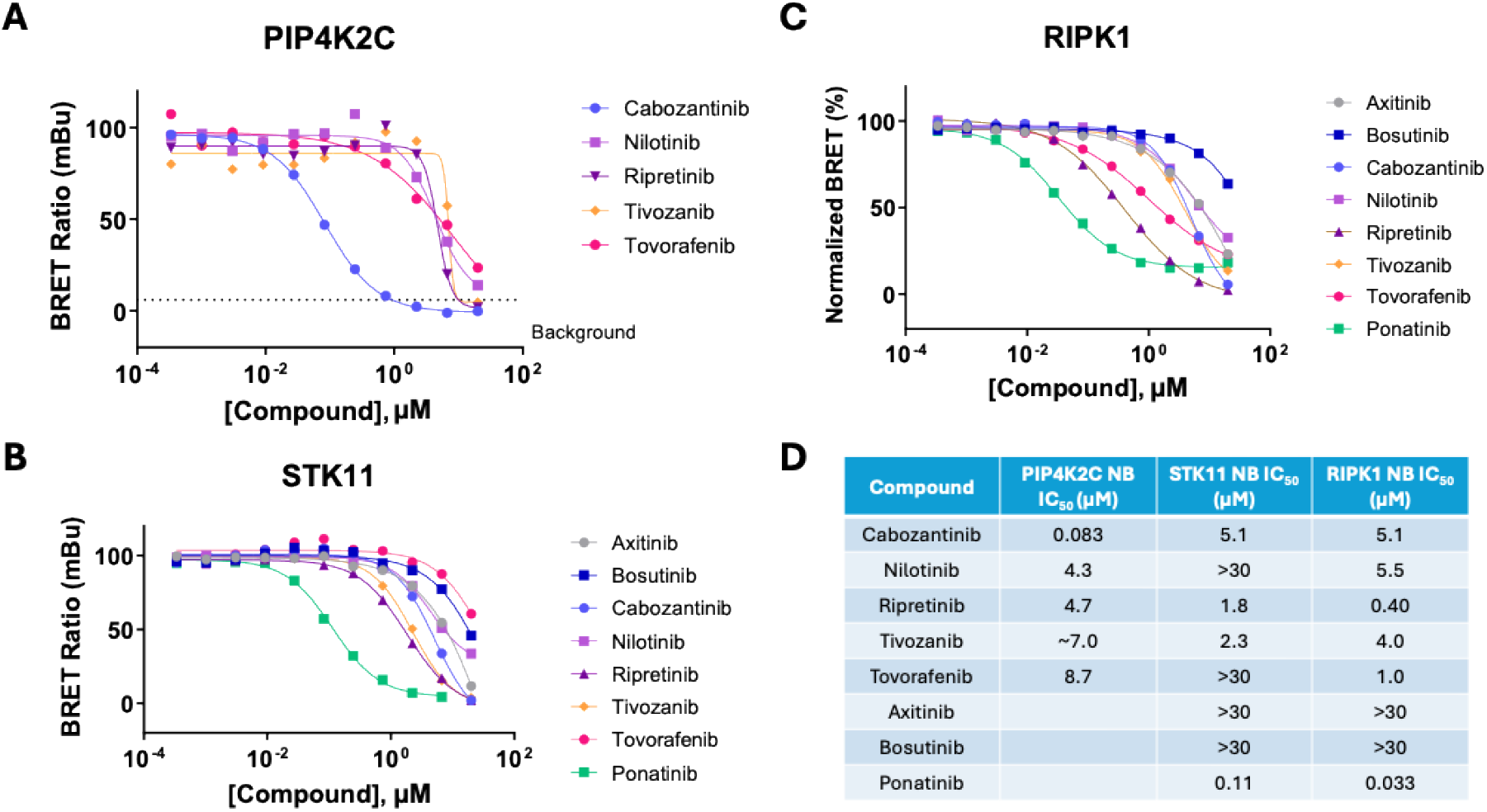
Evaluation of type II inhibitors versus frequent hitter kinases reveals high affinity intracellular interactions for (A) PIP4K2C, (B) STK11, and (C) RIPK1. (D) IC_50_ values from these assays have been tabulated. Each data point represents the mean of two technical replicates.

#### Cabozantinib potently engages PIP4K2C in cells

While PIP4K2C was engaged by several FDA-approved type II inhibitors in the high micromolar range, more potent engagement was observed with the FDA-approved inhibitor cabozantinib (Figure 4A). As an inhibitor of multiple receptor tyrosine kinases, the precise mechanism of action of cabozantinib in the cancers for which it is approved is only partially determined and attributed to activity at MET, AXL, RET, and VEGFR2. The potent engagement of cabozantinib with PIP4K2C is noteworthy, as cabozantinib attains a total C_max_ exceeding its NanoBRET IC_50_ of 83 nM.^34^ Though speculative, this unexpected engagement may underlie critical elements of its mechanism of action as an immune-oncological (I/O) combination therapy. PIP4K2C impacts cytokine release in mice,^35^ a phenomenon perhaps paralleled in the immune-permissive tumor microenvironment seen with cabozantinib combination therapies.^36^

#### Ponatinib potently engages tumor suppressor STK11 (LKB1) in cells

As a well-studied tumor suppressor, pharmacological STK11 blockade is conceptually worrisome. STK11 emerged as a frequent hitter in type II drug profiling in the NanoBRET system, demonstrating engagement of cabozantinib, ripretinib, and tivozanib in the low micromolar range (Figure 4B). Ponatinib engaged STK11 and several other kinases more potently in cellular NanoBRET studies compared to cell-free analysis. Ponatinib demonstrates engagement of STK11 with a NanoBRET IC_50_ = 110 nM, below its total reported clinical C_max_.^37^ Thus, off-target inhibition of the tumor-suppressing function of STK11 may represent a critical liability. Ponatinib shows a uniquely high incidence of toxicities,^38^ and STK11/AMPK suppression in endothelial cells offers a plausible molecular link. These findings position STK11 as a previously unrecognized off-target liability for type II scaffolds and highlight NanoBRET target engagement assays as a critical filter for detecting interactions that may still influence efficacy–toxicity profiles.

#### Tovorafenib potently engages RIPK1 in cells

Developed as a pan-RAF inhibitor, tovorafenib demonstrated low micromolar affinity for RIPK1 in cells (Figure 4C), below its FDA-reported total C_max_.^39^ Since RIPK1 modulates necroptosis and neuroinflammation, this interaction could positively influence anti-tumor immunity. However, such engagement may also impart neuroinflammatory adverse events as a potential liability, meriting orthogonal confirmation and further studies. Although a causal link between such collateral interactions and clinical manifestations remains speculative, live-cell NanoBRET assay profiling unveils a set of hitherto hidden type II interactions that could reshape our understanding of kinase selectivity. Moreover, as conformationally-selective chemotypes, type II kinase inhibitors could uncover unique kinase conformational dynamics present in cells that diverge from those observed in isolated systems.

### A focused screen of select KCGS compounds confirm type II inhibitors as more selective in cells

To further explore the potential type II vulnerability in cells and complement our growing dataset, we sought to compare cell-free and cell-based data for additional type II inhibitors. We selected inhibitors with Eurofins DiscoverX data because it offers the most comprehensive cell-free data and evaluated these compounds in the 240-membered NanoBRET panel. A mix of type I (GW296115X) and type II inhibitors (GW830263A and TPKI-39) were chosen from the kinase chemogenomics set (KCGS).^40^ For KCGS compounds, complete cell-free profiling data had been obtained and published, and K_d_ and/or orthogonal follow-up assays have been executed, providing a compendium of cell-free data for each. Our analyses of the cell-free data for the compounds selected suggested that their selectivity should be variable. Based on the kinome-wide, cell-free profiling data, GW830263A and GW296115X were identified as promiscuous inhibitors, binding to more kinases than TPKI-39.

We show the selectivity fingerprints for these compounds in the Eurofins DiscoverX panel and NanoBRET selectivity panel in Figure 5 when profiled at a common concentration of 1 µM. As was done in Figure 3, data from the cell-free profiling (DiscoverX) was converted to percent inhibition and plotted versus the percent occupancy generated in the cell-based NanoBRET selectivity panel consisting of 240 kinases. Only the wild-type, human kinases that demonstrated >50% binding in both assay formats are plotted in Figure 5 and a correlation line was drawn. Examination of the datasets revealed good agreement of data for kinases in both panels. The biochemical potency of each compound was notably higher when compared to the cell-based assay results. The simplest explanation for this finding is that competition with ATP in the cell reduces the measured affinity of inhibitors. A large portion of cell-free data, however, was not replicated in the cell-based format (data not shown). There was, however, correlative cell-free inhibition data for most of the high affinity binding kinases in cells.

**Figure 5.**
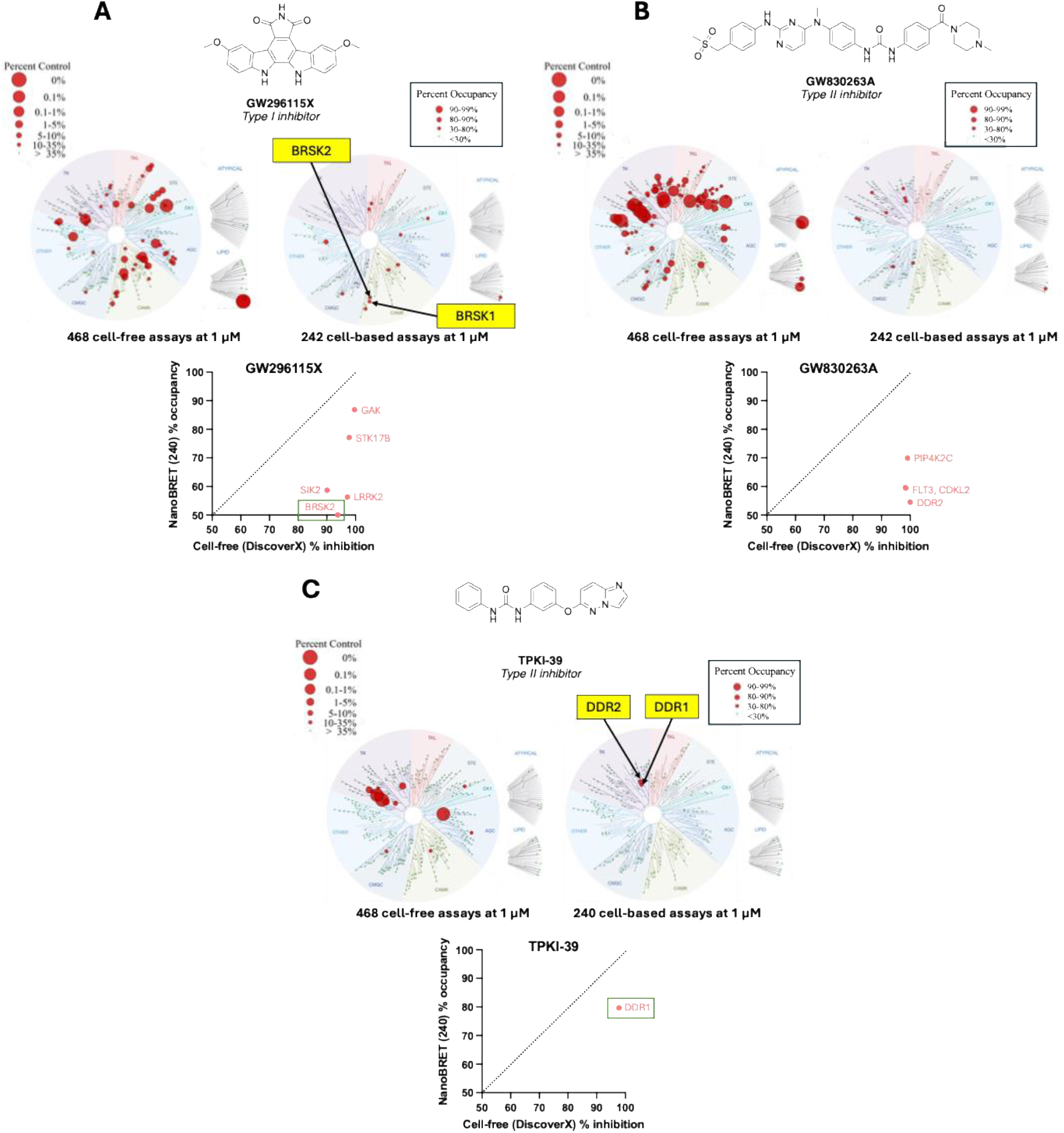
Comparison of percent inhibition and percent occupancy values generated for three kinase inhibitors in cell-free binding and cell-based binding assays, respectively. Data and structures are shown for (A) GW296115X (type I), (B) GW830263A (type II), and (C) TPKI-39 (type II).

#### In-cell profiling reveals GW296115 as best available tool to study BRSK biology

GW296115 has been used as a BRSK2 inhibitor based on our publication, which described it as a cell-active inhibitor of this kinase.^41^ Orthogonal, enzymatic assays were pursued and confirmed potent inhibition (IC_50_ ≤160 nM) of 27 of the 36 highest affinity kinases based on cell-free broad screening. A salt form of this compound, GW296115X, has also been extensively characterized in the DiscoverX cell-free panel.^40^ As expected, these two compounds have similar cell-free profiles in terms of the most high affinity kinases and kinome-wide selectivity scores. The 240-member cell-based panel was used to profile GW296115X, which resulted in a much-improved kinome-wide selectivity profile in cells (Figure 5). Many of the potent cell-free binding kinases did not demonstrate high occupancy in the cell-based screen. BRSK2 was still amongst the high affinity binders in cells, while BRSK1 just missed the in-cell occupancy threshold for inclusion on the graph (42% occupancy). Based upon this in-cell selectivity profile, we assert that GW296115X is amongst the most selective tools to interrogate BRSK biology in cells. Since BRSK1 and BRSK2 have not been extensively characterized, there is great value in identifying chemical tools like GW296115X to accelerate discoveries related to their function.

#### GW830263A is another type II inhibitor that engages PIP4K2C

Like cabozantinib and other type II inhibitors (Figure 4A), GW830263A was found to bind PIP4K2C with high occupancy in cells. This interaction was also identified in the cell-free Eurofins DiscoverX panel. As observed for other type II inhibitors, the in-cell profile of GW830263A identifies fewer interactions than the cell-free screening (Figure 5B). This type II inhibitor represents a chemical starting point that makes high affinity in-cell interactions with kinases from many different families across the kinome and could be further optimized toward a chemical probe for any of these.

### TPKI-39 is confirmed to be an orthogonal DDR chemical probe based on in-cell profiling results

TPKI-39 is a type II inhibitor that demonstrated high affinity binding to 15 kinases with PoC <35 when profiled in the DiscoverX 468-member cell-free panel at 1 µM. Many of these kinases were confirmed to bind when the same assays were run in dose–response format, yielding K_d_ values <100 nM for seven kinases chosen for follow-up analysis. TPKI-39 was also profiled in a chemical proteomics target engagement study using Kinobeads, and DDR2 and PDGFRB were identified as binding to TPKI-39 in cancer cell lysates.^42^ To supplement these data, we selected TPKI-39 for screening in the 240-member NanoBRET selectivity panel. As shown in Figure 5C, TPKI-39 demonstrated occupancy in cells and inhibition in cell-free assays ≥50% for only DDR1 when screened at 1 µM. Of note, DDR2 was identified as a high affinity binder (84% occupancy) in the cell-based assay panel but did not bind (1% inhibition) when profiled in the cell-free assay panel. As a next step, we collated the DiscoverX and 240 cell-based panel single-concentration data and selected a panel of kinases for follow-up in dose–response format (Table 1). Where possible, NanoBRET assays were prioritized for follow-up purposes. If no NanoBRET assay was available, we executed enzymatic assays to confirm cell-free results. Kinases that are present in the K240 screen and did not demonstrate occupancy ≥50% are indicated in red in Table 1. Of the 15 kinases with PoC <35, only 6 were not in the K240 panel (NP in Table 1). There are also 46 kinases in the 240-member panel that are not present in the cell-free assay platform used. As was the case for many other inhibitors, high affinity cell-free data did not replicate when TPKI-39 was profiled in the 240-member in-cell panel (Figure 5C), resulting in a more selective profile for TPKI-39 in cells.

**Table 1.** Combined kinome-wide profiling data and follow-up for TPKI-39.

| Kinase | DiscoverX percent of control | DiscoverX K<sub>d</sub> values (nM) | K240 percent occupancy | NanoBRET IC<sub>50</sub> (nM) | Enzymatic IC<sub>50</sub> (nM) |
| --- | --- | --- | --- | --- | --- |
| CIT | 0 | 1.1 | NP | Low assay window | 3000 |
| KIT | 0 | 0.94 | 19 | >10000 | 11 |
| PDGFRB | 0 | 1.0 | NP | No assay | 1200 |
| CSF1R | 0.4 | 27 | 0 | NB | 230 |
| PDGFRA | 1.1 | 3.6 | NP | No assay | 260 |
| DDR1 | 2.2 | 24 | 80 | 510 | 380 |
| FLT1 | 3.2 | 91 | NP | 480 | 65 |
| TIE2 | 7.6 | NT | 0 | NB | >10000 |
| ABL1 | 11 | NT | 0 | NB | 4700 |
| MAP3K19 | 13 | NT | 4 | NB | 1300 |
| VEGFR2/KDR | 22 | NT | NP | 1500 | 430 |
| CDK8 | 24 | NT | 0 | NB | - |
| PKAC-beta | 30 | NT | 4 | NB | - |
| DCAMKL3 | 31 | NT | 0 | NB | - |
| FLT4 | 33 | NT | NP | No assay | 160 |
| DDR2 | 99 | NT | 84 | 310 | 120 |

NT: not tested; NP: not present; NB: no binding in K240

Based on full-curve NanoBRET assays, TPKI-39 only demonstrates in-cell binding to DDR1, DDR2, and FLT1 (colored green in Table 1) and binds with a similar affinity to all three kinases (Figure S2). As an orthogonal measurement, TPKI-39 inhibits each of these kinases with an enzymatic IC_50_ <400 nM. While robust NanoBRET assays for CIT, PDGFRA, PDGFRB, and FLT4 have not yet been enabled, the enzymatic IC_50_ values imply that high affinity binding of TPKI-39 to these kinases does not result in enzymatic inhibition (CIT and PDGFRB) and/or different protein constructs were used in the respective assays. The NanoBRET IC_50_ value for KIT aligned well with its 19% occupancy value when TPKI-39 was screened broadly in the 240-member in-cell panel at 1 µM. We collected the enzymatic inhibition data for most putative off-target kinases and found good correlation between the enzymatic data and cell-based data for TIE2, ABL1, and MAP3K19, but not for CSF1R. Based on these data, we concluded that TPKI-39 represents a chemical probe for interrogating DDR1/DDR2/FLT1 inhibition in cells. It is worth noting that the cell-free DiscoverX data in isolation suggests high affinity binding of several kinases (CIT, KIT, PDGFRB, PDGFRA, TIE2, ABL1, MAP3K19) and a missed interaction with DDR2. Several of these kinases, including KIT, CSF1R, TIE2, ABL1, and MAP3K19, did not recapitulate this efficacious binding in cells. If the cell-free profiling were considered alone, a high-quality chemical probe would have been overlooked due to its sub-optimal selectivity fingerprint. TPKI-39 provides a tangible example of how the use of in-cell selectivity profiling can guide chemical probe prioritization as well as how it complements rather than replaces cell-free screening.

With a putative chemical probe in hand, we sought a structurally similar analog that lacks DDR1 and DDR2 affinity. Based on broad cell-free screening, TPKI-41^43^ (Figure 6) demonstrated good kinome-wide selectivity (S_35_(1 µM) = 0.047)) and lacked affinity for DDR1 (55 PoC), DDR2 (58 PoC), and FLT1 (21 PoC) when profiled at 1 µM. For reference, a lower PoC corresponds with higher affinity binding, and a lower selectivity score (S_35_(1 µM)) corresponds with higher kinome-wide selectivity. To confirm these data in cells, we executed the respective NanoBRET assays. TPKI-41 was found to be inactive versus DDR1 and DDR2 (IC_50_ >10 µM for both) and exhibit minimal cellular target engagement with off-target kinase FLT1 (4400 nM, Figure S3).

**Figure 6.**
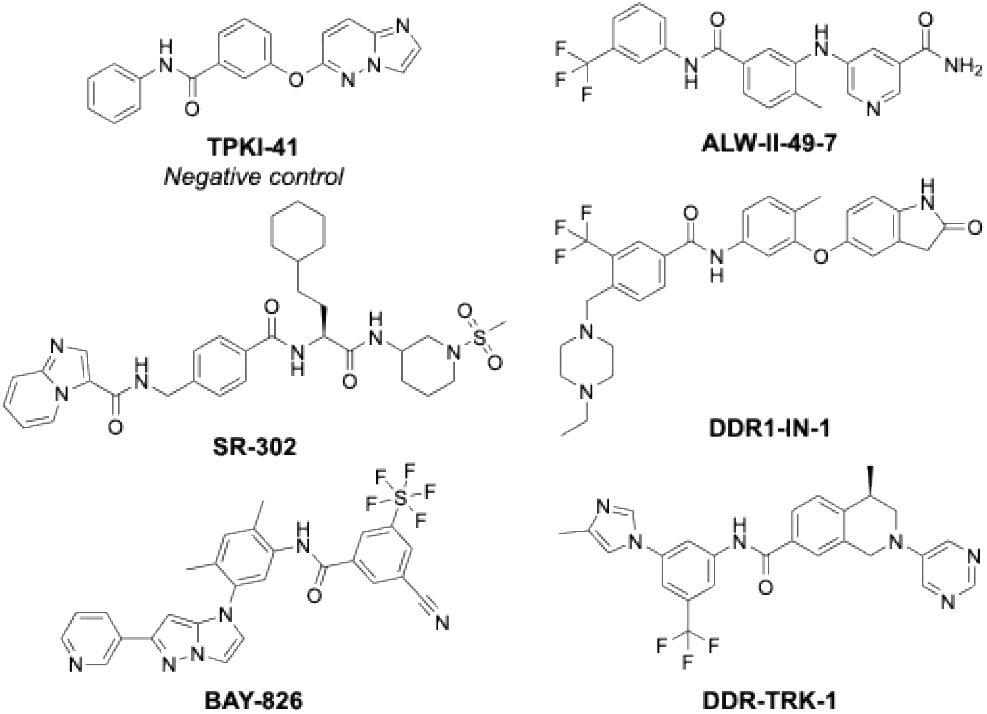
Structure of TPKI-41 and best available DDR1/2 probes.

There are currently five chemical probes listed on the chemical probes portal^44^ for use with DDR1 and DDR2 (Figure 6). Several have also been nominated as DDR probes by the SGC. Reported Eurofins DiscoverX cell-free selectivity profiles for these structurally distinct chemotypes, when profiled at either 100 nM or 1 μM, are listed in Table 2. The S_10_ score of TPKI-39 at 1 µM in the cell-free DiscoverX panel is 0.02, which is comparable to available probe molecules. However, to our knowledge, K192 or K240 in-cell selectivity data have not been collected for these alternative DDR1/2 chemical probes. As shown in Table 2, most DDR chemical probes, like TPKI-39, exhibit potent cellular target engagement of DDR1 and DDR2, with the exception of ALW-II-49-7, which has not yet been evaluated.^45^ DDR1-IN-1 also shows modest selectivity for DDR1 versus DDR2 in the respective NanoBRET assays (Table 2 and Figure S4). The cell-free profiling data for SR-302,^46^ BAY-826,^47^ and DDR1-IN-1^48, 49^ at 100 nM indicate that these inhibitors should be employed at lower concentrations to engage DDR1/2 in cells while minimizing off-target inhibition (Table 2). DDR1-IN-1 and DDR-TRK-1 both have similar reported selectivity S_1_(1 μM) = 0.008 and S_1_(1 μM) = 0.012, respectively.^48-50^ Similar to TPKI-39, most reported DDR1/2 probes also inhibit several off-target kinases. None of the chemical probes reported, however, have overlapping off-target kinases with TPKI-39 at the recommended doses, making it a useful orthogonal probe to those in the literature. TPKI-39 is a potent DDR1/DDR2 chemical probe, with only FLT1 as an off-target kinase at 1 µM in cells. Furthermore, to our knowledge, it is currently the only probe with available broad in-cell selectivity profiling data. Therefore, TPKI-39 is a complementary chemical probe to the current suite of available probes to interrogate DDR1 and 2 biology in cells.

**Table 2.**
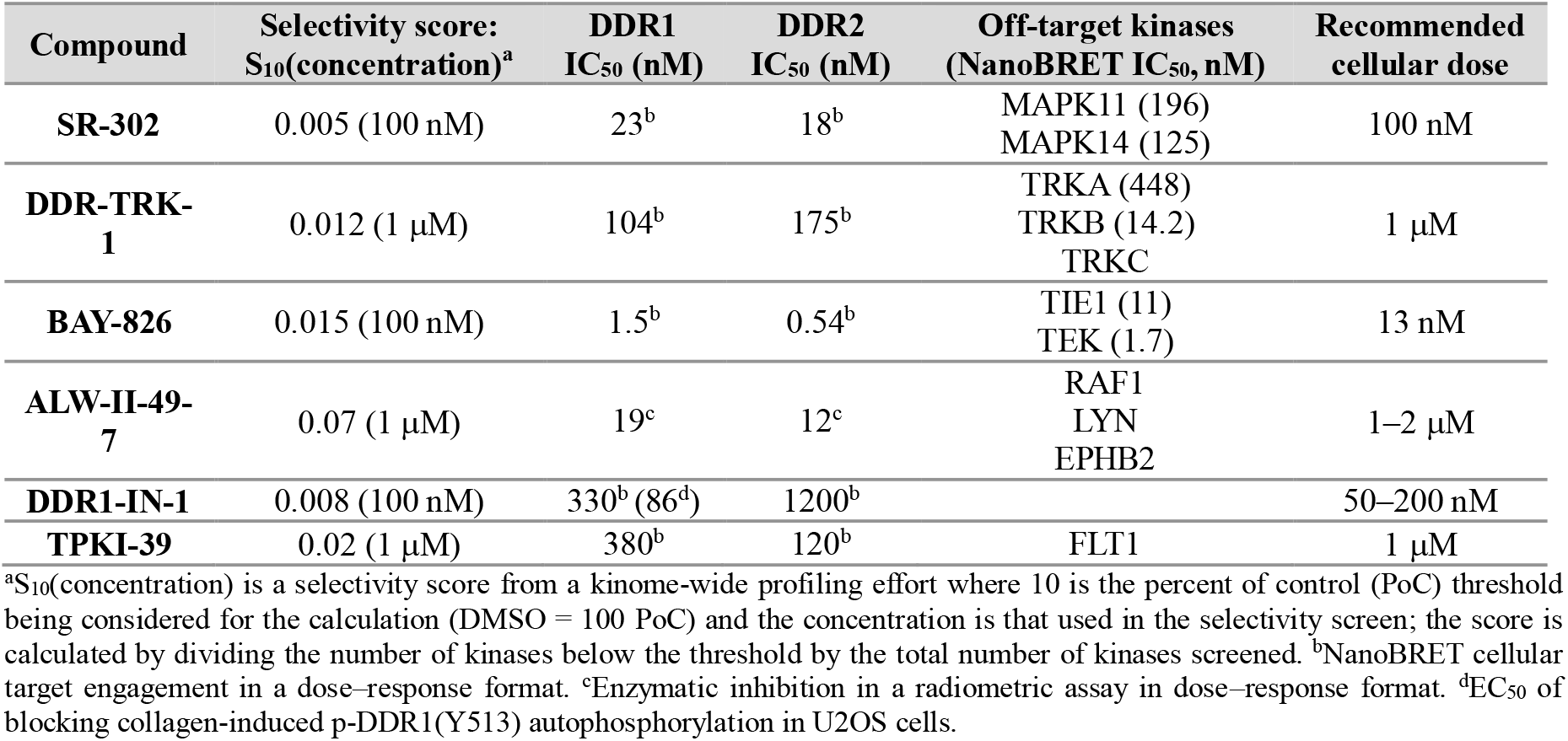
Potency and selectivity data for available DDR1/2 chemical probes.

To confirm inhibition of DDR1 signaling by TPKI-39 in cells, we analyzed DDR1 phosphorylation by western blot. DDR1 has been well-documented to be activated by both fibrillar and basement membrane collagens.^51 52^ Upon collagen binding to the extracellular domain, receptor clustering is induced, facilitating the process of autophosphorylation. Tyrosine residues are phosphorylated in multiple domains of DDR1, including Y792, Y796, and Y797 in the kinase domain,^53^ which are required for activity, as well as Y513 in the intracellular juxtamembrane domain, which is important for binding of downstream protein substrates.^54 55^ Since Y513 is not included in each of the five main DDR1 isoforms, we chose to transiently transfect HeLa cells with a plasmid encoding the canonical DDR1b sequence. As shown in Figures 7 and S5, treatment with TPKI-39 was able to inhibit DDR1 autophosphorylation to basal levels observed without collagen activation in a comparable fashion to established chemical probe DDR1-IN-1.^48, 49^ The negative control, TPKI-41, had no inhibitory effect. These results support the biological activity of TPKI-39 in a cellular setting.

**Figure 7.**
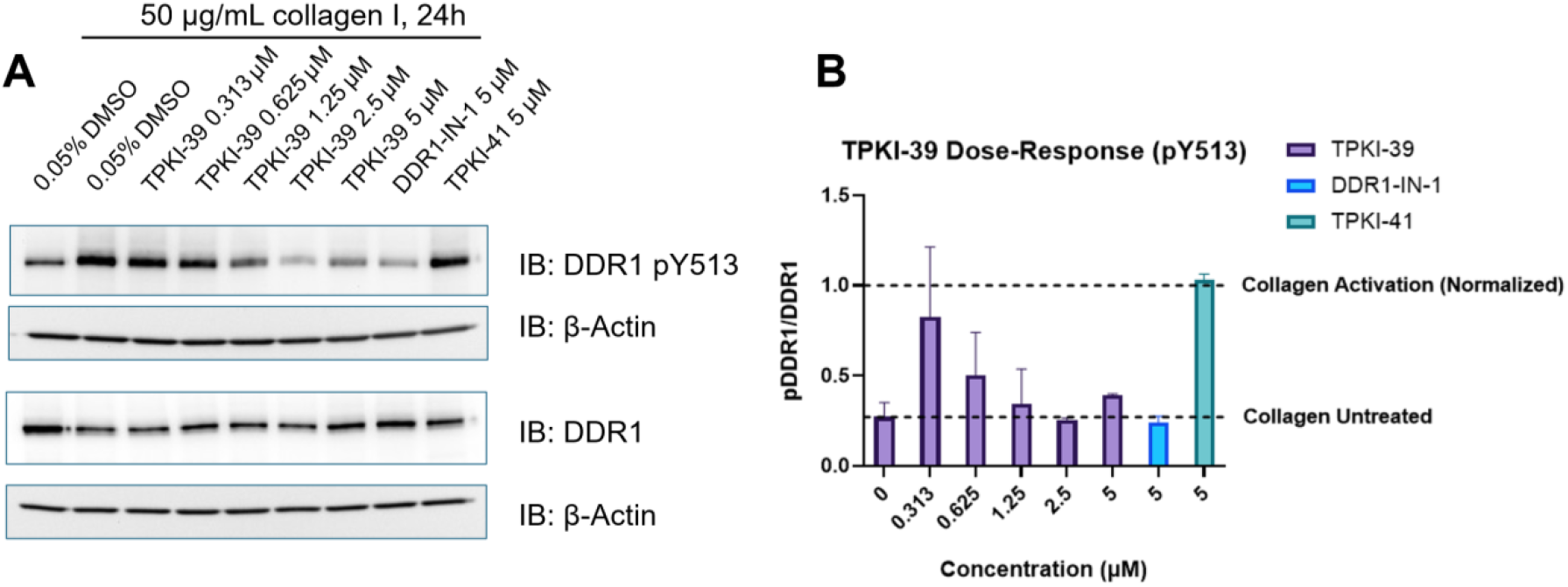
Inhibition of DDR1 autophosphorylation by TPKI-39. (A) Western blotting analysis of TPKI-39 treatment of HeLa cells transiently transfected with cDNA encoding DDR1b isoform. Indicated samples were co-treated with collagen I and TPKI-39. DDR1-IN-1 and TPKI-41 were included as positive and negative controls, respectively. (B) ImageJ quantification of DDR1 western blots. Data is reported as ratio of pDDR1 to total DDR1, normalized to activation by collagen in the absence of inhibitor. Samples are normalized to β-actin loading control. Results are representative of n=3 independent experiments.

## Conclusions

Iteratively benchmarking selectivity is critical during development of small molecule kinase inhibitors. Here, we highlight the power of using cell-based methods, such as NanoBRET cellular target engagement assays, for evaluating selectivity. While current cell-free assay panels are highly sensitive, robust, and comprehensive, in-cell selectivity screening has uncovered high affinity type II interactions that were missed in cell-free profiling. By placing selectivity assays within the context of the cellular environment, biological relevance is increased. Our results support the notion that by interrogating full-length kinases in live cells, unique conformational dynamics can be observed for certain kinase inhibitors that may be less evident with kinase domains in isolation. Additionally, the use of cell-free panels alone may paint an incomplete picture of the kinome-wide selectivity of a lead compound by flagging interactions that are less relevant in a cellular environment. Best practice should involve employing a combination of both cell-free and cell-based assays for the most confident selectivity assessment. TPKI-39 serves as an excellent example of these principles: a chemical probe that may have been overlooked if cell-free selectivity data were considered in isolation. While its cell-free data profile showed engagement of numerous off-target kinases with high affinity, in-cell profiling and follow-up assays illuminated its utility as a potent and selective chemical probe for DDR1/2 and FLT1.

### Experimental Section

#### Compound Characterization

Commercially available compounds used in Figures 2, 3, 4, and 7 and Table 2 were purchased as indicated below.

| Compound | Supplier | Cat No. |
| --- | --- | --- |
| Dasatinib | Selleckchem | S1021 |
| Sorafenib | Selleckchem | S7397 |
| AST487 | Cayman Chemical | 19477 |
| Foretinib | Selleckchem | S1111 |
| LXH254 | Selleckchem | S8745 |
| AMG925 | Selleckchem | S7533 |
| Ponatinib | Selleckchem | S1490 |
| Bosutinib | Selleckchem | S1014 |
| Cabozantinib | MedChemExpress | HY-13016 |
| Nilotinib | Selleckchem | S1033 |
| Axitinib | MedChemExpress | HY-10065 |
| Ripretinib | MedChemExpress | HY-112306 |
| Tivozanib | MedChemExpress | HY-10977 |
| Tovorafenib | MedChemExpress | HY-15246 |
| ALW-II-49-7 | MedChemExpress | HY-18833 |
| DDR1-IN-1 | MedChemExpress | HY-13979 |

GW296115X, GW830263A, TPKI-39, and TPKI-41, which were used in Figures 5 and 7, are part of KCGS,^56^ PKIS,^57^ and/or PKIS2.^40^ DMSO stock solutions of these compounds are housed at and were provided by UNC. DMSO stock solutions of BAY-826, SR-302, and DDR-TRK-1 (Table 2) were provided by the SGC as part of their donated chemical probes program. All compounds used were confirmed to be >95% pure by HPLC analysis.

#### Cell Culture

Human embryonic kidney (HEK293) cells (hypotriploid, female, fetal) were obtained from ATCC (CRL-1573). HEK293 cells were cultured in Dulbecco’s Modified Eagle’s medium (DMEM, Gibco, #11965092) supplemented with 10% fetal bovine serum (FBS). HEK293 cells were incubated at 37 °C in 5% CO_2_ and passaged every 72 hours with 0.05% trypsin (Gibco, #25300054) not allowing them to reach confluency. HeLa cells were obtained from ATCC (CCL-2) and cultured in DMEM (Gibco, #11965092) supplemented with 10% FBS. HeLa cells were incubated at 37 °C in 5% CO_2_ and passaged every 72 hours with 0.05% trypsin (Gibco, #25300054) not allowing them to reach confluency.

#### NanoBRET Assays

Kinase constructs used with associated tracer concentrations: KIT-NLuc (0.063 µM K-4), KDR-NLuc (0.5 µM K-9), DDR1-NLuc (0.031 µM K-4), DDR2-NLuc (0.063 µM K-4), FLT1-NLuc (0.33 µM K-9), PIP4K2C-NLuc (0.063 µM K-8), NLuc-STK11 (0.5 µM K-10), and NLuc-RIPK1 (0.33 µM K-9). NanoBRET assays were performed as previously described.^15, 58^ HEK293 cells were transfected with NLuc tagged kinase constructs as previously described, and 100 µL of HEK293 cells in DMEM + 10% FBS were plated in 96-well plates (Corning, #3917) at a cell density of 2 × 10^5^ cells/mL. After 16-20 hours, HEK293 complete media was replaced with Opti-MEM without phenol red, with 100 µL in no tracer wells, 95 µL in tracer only wells, and 85 µL in tracer and compound wells. Plates were incubated with compound at 37 °C in 5% CO_2_ for 2 hours. BRET was generated as recommended in the manufacturer’s protocol (50 µL of a 3X solution of NanoBRET NanoGlo substrate and extracellular inhibitor). Raw milliBRET units (mBU) were read on a GloMax Discover (Promega) with a donor emission wavelength (450 nm) and an acceptor wavelength (600 nm). mBU were calculated by dividing the acceptor values by the donor values, subtracting an average of the no tracer containing wells, and multiplying by 1,000. Log(inhibitor) vs response (three parameter) dose–response curves in mBU were normalized in GraphPad and used to generate an IC_50_.

#### Enzymatic Assays

Radiometric assays were performed at the Km value for ATP of the specific kinase at Reaction Biology Corporation (RBC). Compounds were tested in a 10-point dose–response format to generate the IC_50_. Details of the assay protocol, including substrate, protein constructs, and controls are available at the RBC website: https://www.reactionbiology.com/list-kinase-targets-us-facility.

#### Selectivity Profiling Using K240 and K300 Assays

Kinase selectivity profiling in the K240 and K300 NanoBRET panels was performed by Promega’s Tailored R&D Solutions (TRS) Screening Services team. TRS has optimized and adopted NanoBRET TE for high-throughput compound selectivity profiling of kinase inhibitors.^59^ Each panel was run in technical duplicate and the average values obtained plus standard deviation are displayed in the respective SI file. The choice of tracers and appropriate concentrations for the K300 NanoBRET panel, which are the same as used in the K240 NanoBRET panel, have also been included as an SI file.

#### Western Blotting

HeLa cells were seeded overnight at 300-350K cells/well and transiently transfected with cDNA encoding DDR1b isoform for 20-24 hours. Media was replaced with fresh media containing the indicated concentrations of DMSO, TPKI-39, DDR1-IN-1, or TPKI-41. Cells were pre-treated for 1 hour before media was exchanged for media containing the same compound concentration as well as 50 ug/mL rat tail collagen I (Enzo, ALX-522-435-0020) and treated for an additional 24 hours. Cells were washed with PBS and scraped in 1X RIPA buffer supplemented with Halt protease/phosphatase inhibitor cocktail (Thermo). 20 ug total protein was run on 4-12% tris-glycine gels (Thermo) and transferred to PVDF membranes. Phospho-blots were blocked in 5% BSA in TBST for 2 hours, washed 3 x 5 minutes with TBST and incubated overnight with mouse β-actin and rabbit DDR1 antibodies (CST). Total DDR1 blots were blocked in 5% dry milk in TBST for 2 hours, washed 3 x 5 minutes with TBST and incubated overnight with mouse β-actin and rabbit DDR1 antibody (CST). The following day, blots were washed 3 x 10 minutes with TBST, incubated 90 minutes with fluorescent (LICOR) or HRP-linked (Invitrogen) secondary antibodies and washed 3 x 10 minutes. Blots were imaged using iBright (Thermo) and quantified with ImageJ.

## Supporting information

Cell-Based Profiling Data

Cell-free Profiling Data

## Associated Content

### Supporting Information

The Supporting Information is available free of charge at http://pubs.acs.org.

Molecular Formula Strings file (CSV); Compound structures, NanoBRET curves, and biological replicates of TPKI-39 western blots (PDF); Percent occupancy data (XLSX); Cell-free profiling data (XLSX); K300 tracer conditions (CSV) are included.

## Author Contributions

#M.B. and F.M.B. contributed equally to this work. All authors have given approval to the final version of the manuscript.

## Notes

K.D.H., M.B.R., C.D., and M.R.S. are employees of Promega. Promega owns patents related to NanoBRET technology. The remaining authors declare no competing financial interest.

## Acknowledgment

Kinase constructs for NanoBRET assays run at UNC were kindly provided by Promega with NLuc tag orientations listed: KIT-NLuc, KDR-NLuc, DDR1-NLuc, DDR2-NLuc, and FLT1-NLuc. Coral was used to make the kinome illustrations in the Table of Contents graphic. Coral was developed in the Phanstiel Lab at UNC; https://phanstiel-lab.med.unc.edu/CORAL.^60^

The SGC is a registered charity (number 1097737) that receives funds from Bayer AG, Boehringer Ingelheim, the Canada Foundation for Innovation, Eshelman Institute, Genentech, Genome Canada through Ontario Genomics Institute, EU/EFPIA/OICR/McGill/KTH/Diamond, Innovative Medicines Initiative 2 Joint Undertaking, Janssen, Merck KGaA (aka EMD in Canada and USA), Pfizer, the São Paulo Research Foundation-FAPESP, and Takeda. This work was supported in part by an NIH T32 Chemistry-Biology Interface training grant (5T32GM135122).

## Abbreviations

AGC: protein kinase A, G, C
ATP: adenosine triphosphate
c-Abl: Abelson tyrosine-protein kinase 1
CAMK: calcium/calmodulin-dependent protein kinases
cDNA: complementary DNA
CK1: casein kinase 1
C_max_: maximum concentration
CMGC: Cyclin-dependent kinase, mitogen-activated protein kinase, glycogen synthase kinase, CDC2-like kinase
DFG: Aspartic acid-phenylalanine-glycine
DMSO: dimethyl sulfoxide
DNA: deoxyribonucleic acid
FDA: Food and Drug Administration
IC_50_: half maximal inhibitory concentration
KCGS: kinase chemogenomics set
K_d_: equilibrium dissociation constant
K_m_: Michaelis constant
LINCS: Library of Integrated Network-based Cellular Signatures
NanoBRET: bioluminescence resonance energy transfer using NanoLuc
NLuc, nM: nanomolar
pDDR1: phospho-DDR1
PoC: Percent of Control
SGC: Structural Genomics Consortium
STE: sterile kinase
TK: tyrosine kinase
TKL: tyrosine kinase-like
μM: micromolar.

