## Supplemental Figures S1-S5 for "Cellular Context Influences Kinase Inhibitor Selectivity"

### Table of Contents

Figure S1 S2

Figure S2 S2

Figure S3 S3

Figure S4 S3

Figure S5 S4

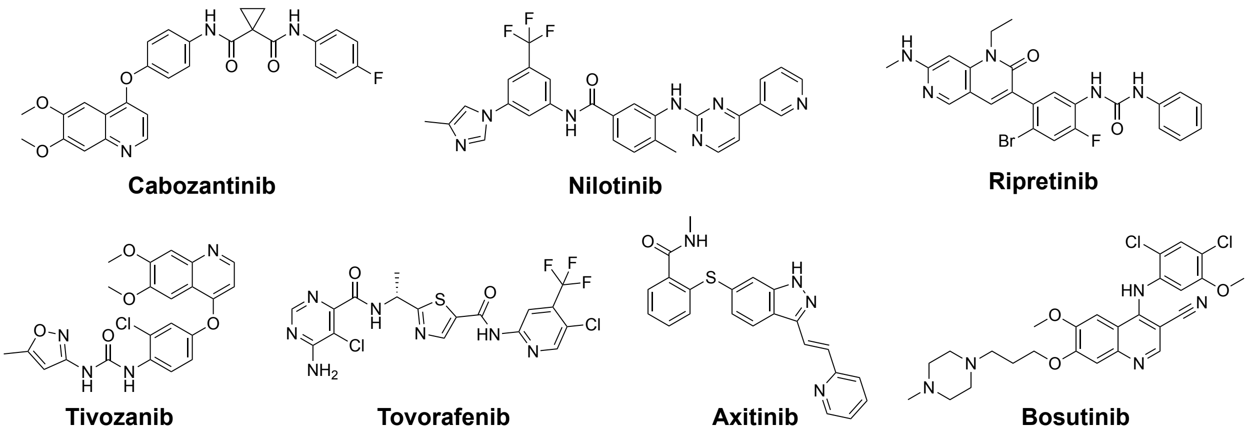


**Figure S1.** Structures of the type II kinase inhibitors screened versus kinases PIP4K2C, RIPK1, and STK11 in NanoBRET target engagement assays.

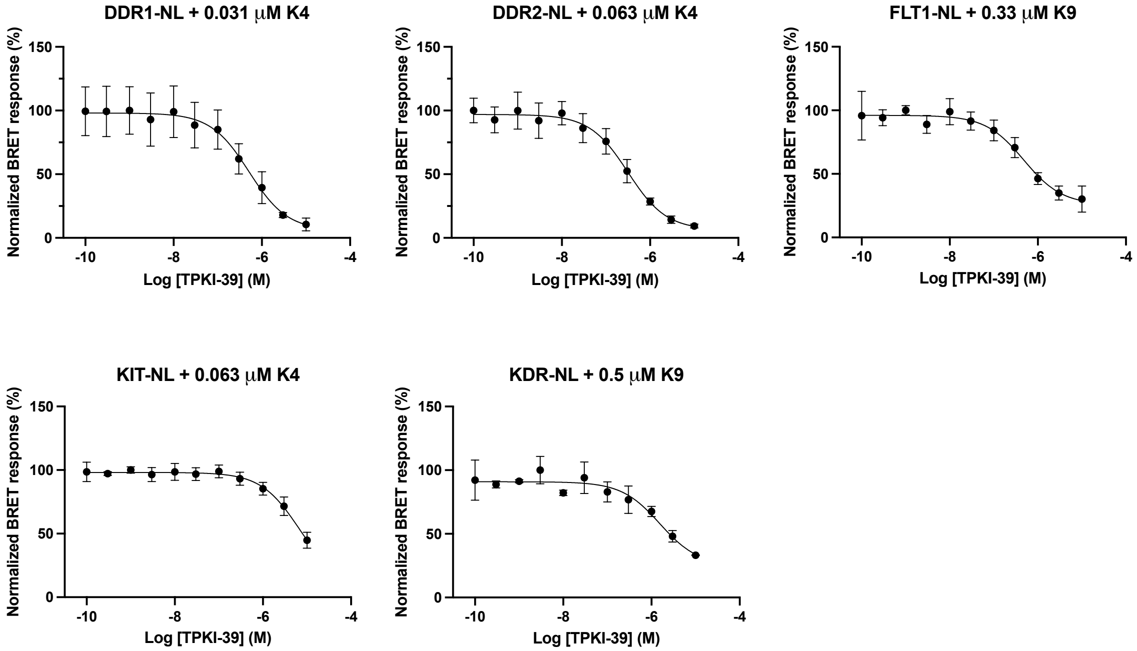


**Figure S2.** NanoBRET curves corresponding to K192/240 follow-up assays for TPKI-39. NanoBRET assays were run with one biological replicate with two technical replicates each (n=1) for KIT and KDR, and in biological duplicates with two biological replicates each (n=2) for DDR1, DDR2, and FLT1. Data were plotted in GraphPad Prism using a log(inhibitor) vs. response (three parameters), and error bars represent the standard error of the mean (SEM).

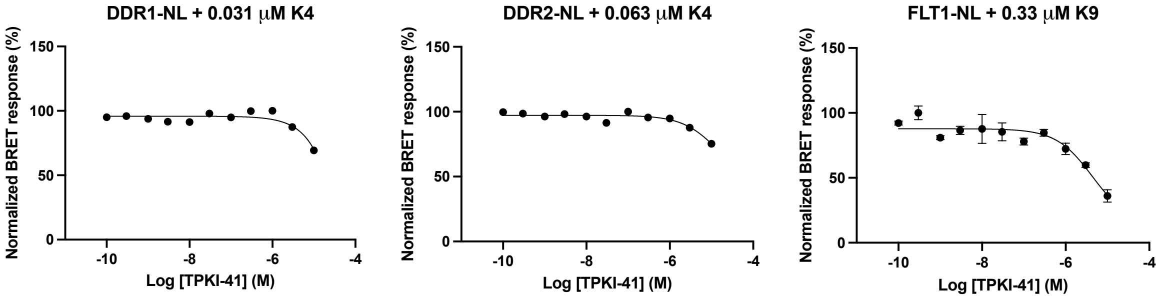


**Figure S3.** NanoBRET curves corresponding for TPKI-41. NanoBRET assays were run with one biological replicate with two technical replicates each (n=1) for FLT1, and with one technical replicate each (n=1) for DDR1 and DDR2. Data were plotted in GraphPad Prism using a log(inhibitor) vs. response (three parameters), and error bars represent the standard error of the mean (SEM).

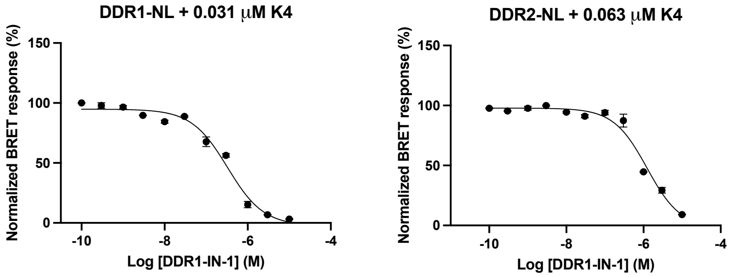


**Figure S4.** NanoBRET curves for DDR1-IN-1. NanoBRET assays were run with one biological replicate with two technical replicates each (n=1). Data were plotted in GraphPad Prism using a log(inhibitor) vs. response (three parameters), and error bars represent the standard error of the mean (SEM).

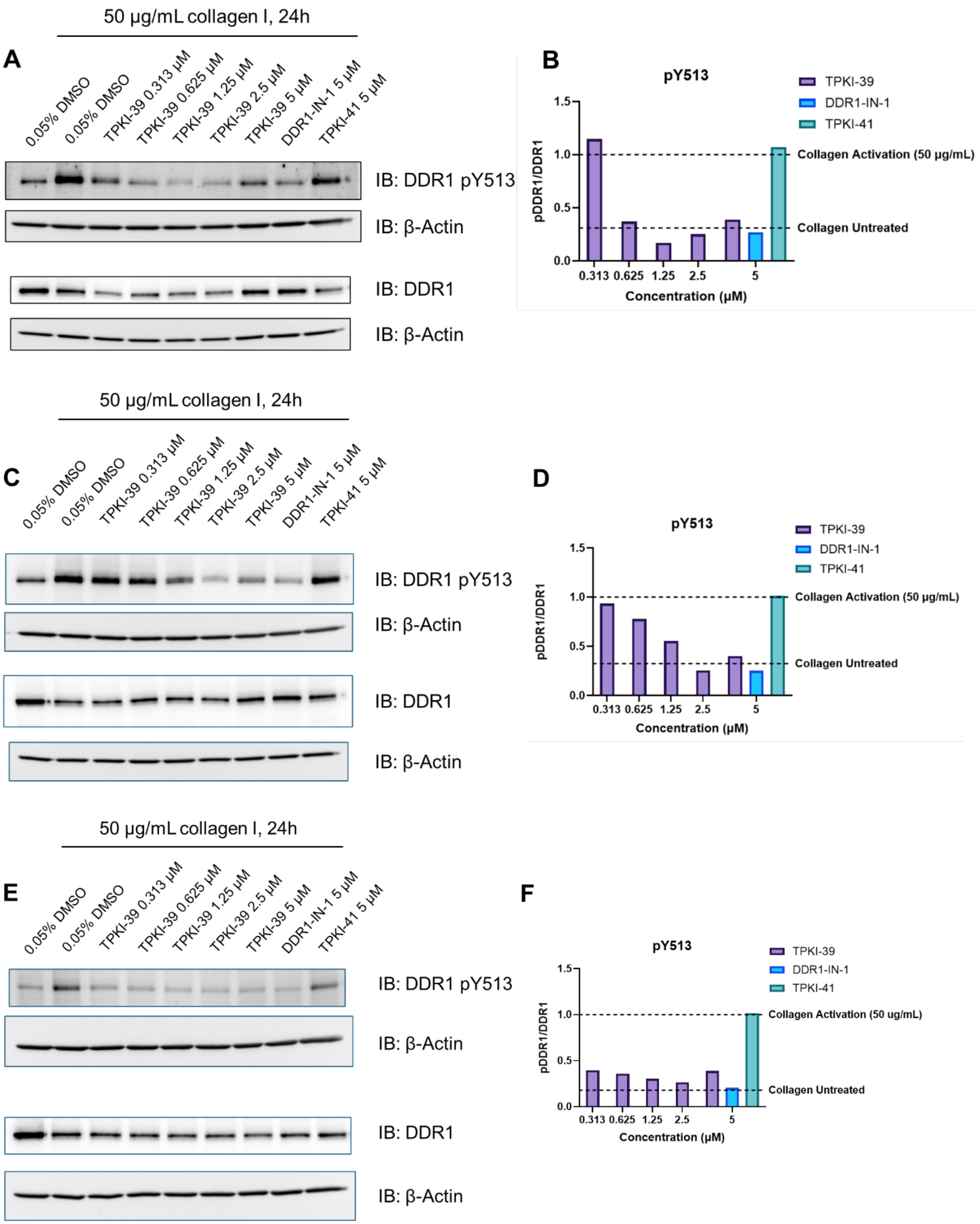


**Figure S5.** Three biological replicates of TPKI-39 inhibition. (A,C,E) Western blotting analysis of TPKI-39 treatment of HeLa cells transiently transfected with cDNA encoding DDR1b isoform. Indicated samples were co-treated with collagen I and TPKI-39. DDR1-IN-1 and TPKI-41 were included as positive and negative controls, respectively. (B, D, F) ImageJ quantification of respective western blots. Data is reported as ratio of pDDR1 to total DDR1, normalized to activation by collagen in the absence of inhibitor. Samples are normalized to β-actin loading control.
